# Novel protein interaction network of human calcitonin receptor-like receptor revealed by label-free quantitative proteomics

**DOI:** 10.1101/2023.04.18.537143

**Authors:** Dimitrios Manolis, Shirin Hasan, Camille Ettelaie, Anthony Maraveyas, Darragh P. O’Brien, Benedikt M. Kessler, Holger Kramer, Leonid L. Nikitenko

## Abstract

**Background:** G protein-coupled receptor (GPCR) calcitonin receptor-like receptor (CLR) signalling is implicated in skin-related and cardiovascular diseases, migraine and cancer. However, beyond its agonists and receptor activity-modifying proteins (RAMPs), proteins which bind to CLR and define its properties in primary human cells remain insufficiently understood.

**Aim:** We aimed to profile the CLR interactome in primary human dermal lymphatic endothelial cells (HDLEC), where this GPCR is expressed.

**Materials and methods:** Immunoprecipitation (IP) of core- and terminally-glycosylated CLR from primary *in vitro* cultured HDLEC was conducted using rabbit polyclonal anti-human CLR serum (with pre- immune serum serving as a control) and confirmed by immunoblotting. Total HDLEC and co-immunoprecipitated CLR proteomes were analysed by label-free quantitative nano-liquid chromatography-tandem mass spectrometry (nLC-MS/MS). Quantitative *in-situ* proximity ligation assay (PLA) using ZEISS LSM 710 confocal microscope and ZEN Blue 3.0 and Image J software was performed to confirm nLC-MS/MS findings. All experiments were repeated at least three times (biological replicates). For statistical analysis of PLA data, distribution was analysed using Shapiro-Wilk normality test followed by an unpaired *t*-test or Mann-Whitney test with a *p*-value of ≤0.05 interpreted as significant. For MS data of CLR IP samples, statistical analysis was performed using *t*-test with a permutation-based false discovery rate (FDR)-adjusted *p*-value of ≤0.006 interpreted as significant.

**Results:** A total of 4,902 proteins were identified and quantified by nLC-MS/MS in primary HDLEC and 46 were co-immunoprecipitated with CLR (*p<*0.006). Direct interaction with the GPCR was confirmed for five of these by PLA (*p*<0.01).

**Conclusions:** This is the first study of its kind to identify novel binding partners of CLR expressed in primary human cells. Our integrative quantitative approach, combining immunoprecipitation of core- and terminally-glycosylated CLR, nLC-MS/MS, and PLA, could be applied in a similar fashion to study its interactome in a variety of human cells and tissues, and its contribution to a range of diseases, where the role of this GPCR is implicated.

## Introduction

Skin dermis is rich in dermal lymphatic vessels and their dysfunction is implicated in skin- related diseases, including lymphoedema and cancer (Carlson, 2014; Petrova & Koh, 2018; Oliver et al., 2020). Lymphatic vessels are lined by highly differentiated lymphatic endothelial cells (LEC), which express specific markers, including podoplanin (PDPN), vascular endothelial growth factor receptor 3 (VEGFR3) and lymphatic vessel endothelial hyaluronan receptor 1 (LYVE1). LEC differentiation and function is regulated by cell surface receptors, including receptor tyrosine kinases (RTKs), G protein-coupled receptors (GPCRs) and others (Zheng et al., 2014).

The calcitonin receptor-like receptor (CLR) is a class B (secretin receptor family) GPCR, encoded by *CALCRL* gene (Flühmann et al., 1995). CLR is expressed at mRNA and protein levels in lymphatic vessels of human skin tissue and in isolated primary human dermal microvascular, including lymphatic, EC (HDMEC and HDLEC respectively) cultured *in vitro* (Nikitenko et al., 2003; Nikitenko et al., 2006; Jin et al., 2008; Maybin et al., 2011; Nikitenko, Shimosawa et al., 2013). In primary HDLEC, CLR regulates proliferation, migration and monolayer permeability (Fritz-Six et al., 2008; Jin et al., 2008; Chen et al., 2011; Davis et al., 2017). In humans, loss-of-function mutation (in-frame deletion of V205) of *CALCRL* is lethal due to non-immune hydrops fetalis during embryogenesis (Mackie et al., 2018). In murine skin, total conditional knockout of *Calcrl* leads to developmentally arrested lymphatic capillaries, dermal lymphangiectasia, increased lymphatic permeability and prolonged oedema (Hoopes et al., 2012). The roles for CLR function in other than skin tissues, and its implication in diseases and disorders such as clear cell renal cell carcinoma (CCRCC), acute myeloid leukaemia (AML), stroke and migraine have been reported (Angenendt et al., 2019; Edvinsson et al., 2018; Gluexam et al., 2019; Herlambang et al., 2021; Koyama et al., 2017; Larrue et al., 2021; Nikitenko, Leek, et al., 2013). The use of monoclonal antibody-based drugs targeting CLR-mediated function for migraine prophylaxis, such as erenumab and galcanezumab, has been associated with impaired wound healing in skin, along with development of skin reactions and side effects such as swelling, rashes or pruritus (Bangs et al., 2020; Wurthmann et al., 2020; Göbel et al., 2022; Schenk et al., 2022). Altogether, these findings warrant investigation of CLR properties in HDLEC and other human cell types where it is expressed in health and disease.

The glycosylation state and transport of CLR from endoplasmic reticulum (ER) to the plasma membrane is determined by heterodimerisation of the GPCR with receptor activity-modifying proteins (RAMP) 1, 2 and 3 (McLatchie et al., 1998; Hilairet, Foord, et al., 2001). The localisation of core glycosylated CLR (∼45 kDa) has been associated with ER, while the mature terminally glycosylated form (∼55 kDa) with receptor expression on the surface of the cells, including primary HDMEC (McLatchie et al., 1998; Nikitenko et al., 2006). Three peptide hormones act as putative agonists of CLR - adrenomedullin (AM), calcitonin gene related peptide (CGRP), and adrenomedullin2/intermedin (ADM2 or IMD) (Poyner et al., 2002). These agonists bind CLR with differential affinity, which is determined by specific RAMPs: CGRP has higher affinity for CLR/RAMP1, while AM and IMD have preference for CLR/RAMP2 and CLR/RAMP3 complexes, respectively, when using *in-vitro* models with ectopic expression of HA, Myc or GFP tagged CLR-RAMP complexes (McLatchie et al., 1998; Poyner et al., 2002; Hay et al., 2018). RAMPs and CLR agonists) also selectively influence trafficking of this GPCR (Kuwasako et al., 2000; Bomberger, Parameswaran, et al., 2005; Cottrell et al., 2007). In human embryonic kidney (HEK293T) cells ectopically expressing individual CLR/RAMP complexes, CLR/RAMP1 heterodimers were preferentially internalised in response to stimulation with CGRP, whilst CLR/RAMP2 and CLR/RAMP3 with AM (Kuwasako et al., 2000). Furthermore, in the presence of N-ethylmaleimide-sensitive factor (NSF) and Na+/H+ exchanger regulatory factor-1 (NHERF-1), RAMP3 (in contrast to RAMP1 and RAMP2) alter CLR trafficking to a recycling pathway after agonist-induced endocytosis or inhibit CLR/RAMP3 complex internalisation respectively (Bomberger, Parameswaran, et al., 2005; Bomberger, Spielman, et al., 2005).

Studies of CLR interaction with other proteins (beyond RAMPs and agonists), have been conducted using immunoprecipitation of ectopically expressed, including tagged CLR/RAMP complexes (Evans et al., 2000; Hilairet, Bélanger, et al., 2001; Padilla et al., 2007). These reports demonstrated that CLR/RAMP heterodimers can interact (either directly or indirectly) with molecules from several classes, including enzymes (e.g. endothelin converting enzyme 1, ECE1), accessory (e.g. receptor component protein, RCP) and scaffold (e.g. β-arrestin2) proteins. However, the limitations of transfected systems to study GPCR pharmacology which is influenced by cell-specific factors have been acknowledged (Hay et al., 2018). In particular, there are inconsistencies between endogenous and transfected models of CLR antagonism, in which the function and regulation of this GPCR were studied. Therefore, the use of primary human cells endogenously expressing (i.e. non-overexpressed or knocked-down) CLR species, such as HDLEC and others (Nikitenko et al., 2006; Jin et al., 2008; Fritz-Six et al., 2008; Nikitenko et al., 2013; Davis et al., 2017; Seyedabadi et al., 2019), provide better models for studying properties, function and regulation of GPCR in human tissues. In particular, the discovery of proteins interacting with CLR (herein termed “CLR interactome”) has the potential to unravel the pharmacology and previously unknown properties of CLR. This knowledge could promote translational research related to diagnostic and prognostic utility of this receptor in health and disease. To our knowledge, there are no data regarding the interactome of endogenous core- and terminally- glycosylated CLR interactome in primary human cells and tissues.

Quantitative MS-based approaches, including label-free proteomics, have been successfully used to discover novel protein-protein interactions (Yugandhar et al., 2019). Immunoprecipitation of endogenously expressed receptor using antibodies in conjunction with MS has been effectively utilised for the identification of GPCR-associated protein complexes, including β2-adrenergic receptor (β2-AR), protease activated receptor 2 and parkin-associated endothelin-like receptor (Chung et al., 2013; Sokolina et al., 2017). Proximity ligation assay (PLA) has emerged as a powerful tool for detection and quantification of protein-protein interactions *in-situ*, along with validation of MS data (Söderberg et al., 2008; Weibrecht et al., 2010). The aim of the present study was to identify binding partners of CLR in *in vitro* cultured HDLEC by using label-free quantitative MS-based proteomics. Herein, we used a combination of immunoprecipitation, nLC-MS/MS and *in-situ* PLA analyses to discover a group of 46 novel proteins interacting with CLR endogenously expressed in primary HDLEC. Our study paves the way for new opportunities and avenues to dissect the CLR interactomes that define the properties and function of this GPCR in a range of human cells and tissues.

## Results

To investigate endogenous CLR interactome in HDLEC, we used an MS-based label-free quantitative approach which also combined non-MS-based methodologies (Figure 1A). Characterisation of HDLEC using immunofluorescence (IF) demonstrated that a pure population of HDLEC (Figure 1B) express CLR, which is localised in both the cell surface and perinuclear space (Figure 1C) Immunoblotting analysis revealed the expression of both core- and terminally-glycosylated CLR in the lysates of *in vitro* cultured HDLEC (Figure 1D). Label-free nLC-MS/MS analysis of HDLEC lysate enabled the identification and quantification of CLR (a total of 2 peptides identified, covering 6.7% of the protein sequence) amongst a comprehensive proteomic profile consisting of 4,902 protein groups, following removal of 99 known contaminating species (Figure 1E, F). The relative abundance of CLR compared to selected lymphatic and pan-endothelial markers, such as von Willebrand factor (vWF), cadherin 5 (CDH5) and platelet and endothelial cell adhesion molecule 1 (PECAM1) (Figure 1F). Gene Ontology cellular component (complete) and Protein Analysis Through Evolutionary Relationships (PANTHER) analyses mapped the predicted distribution (in percentage) of the HDLEC proteome in terms of their protein classes (Figure 1G), along with their subcellular localisation (Figure 1H), to which potential CLR interacting partners could belong. Protein classes included enzymes (33%), transcriptional/translational regulators (26%), cytoskeletal proteins (12%) and signal transducers/modulators (10%) amongst others (Figure 1G). Proteind expressed in HDLEC are mainly associated with the nucleus (33%), plasma membrane (16.5%), mitochondrion (14%), cytoskeleton (11%), endoplasmic reticulum (10%) and (10%), and endosomal-lysosomal system (10%) (Figure 1H).

**Figure 1.**
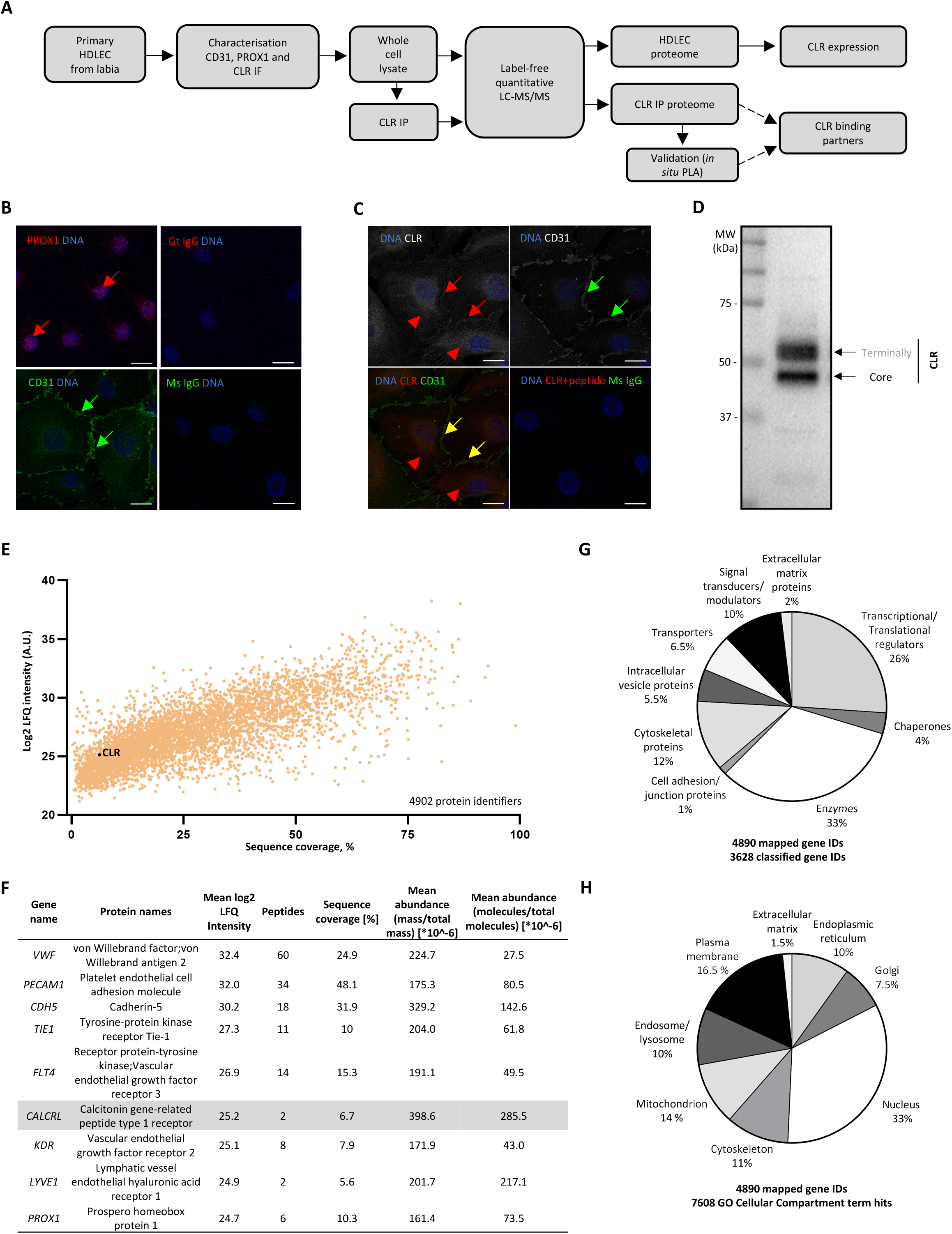
CLR expression in the context of proteome profile of *in vitro* cultured human dermal lymphatic endothelial cells. (**A**) Flow diagram of experimental design of the study. Primary human dermal lymphatic endothelial cells (HDLEC) were cultured *in vitro* and characterised for expression of pan-endothelial cell (pan-EC) and lymphatic endothelial cell (LEC) markers (CD31 and PROX1 respectively) and calcitonin receptor-like receptor (CLR) by immunofluorescence (IF). Anti-human CLR serum/polyclonal antibody captured with protein G magnetic beads was used for CLR immunoprecipitation (IP). Quantitative profiling of HDLEC and CLR co-IP proteome was carried out upon trypsin digestion followed by label-free nano-liquid chromatography-tandem mass spectrometry (nLC-MS/MS) and data- dependent acquisition (DDA) mode. Validation of identified by MS CLR interacting partners was assessed by *in-situ* proximity ligation assay (PLA). Acquired MS data for whole cell lysate and CLR IP were submitted for database searching, quantification, and Gene Ontology (GO) enrichment analysis. For full details, please see Materials and Methods (Supplemental file 1). (**B-C**) Immunofluorescence was performed using primary antibodies and secondary Alexa 488- and Alexa 594- conjugated antibodies (green and red signals respectively). (**B**) HDLEC characterisation using anti-CD31 (green colour) and anti-PROX1 (red colour) antibodies. PROX1 expression in the nucleus (red arrows) and CD31 expression at cell-cell contacts (green arrows) are indicated. Nuclei were counterstained with DAPI (blue colour). (**C**) Subcellular localisation of endogenous CLR in HDLEC detected by IF. CLR is expressed on plasma membrane (red arrows) upon colocalisation with CD31 (yellow colour; yellow arrows) and intracellularly in the perinuclear space (red arrowheads). (**B, C**) Scale bars represent 20 μm. (**D**) Immunoblotting analysis of core- and terminally-glycosylated CLR endogenous expressed in HDLEC. (**E-G**) Label-free quantitative nLC-MS/MS analysis of HDLEC lysates (n=4). (**E**) Scatter plot of HDLEC lysates, in which the percentage of amino acid sequence coverage is plotted against the log_2_ label-free quantitation (LFQ) intensity. Each dot in the scatterplot represents a quantified protein and CLR (encoded by *CALCRL* gene) is highlighted in black. In summary, 56,930 peptides and 5,102 proteins groups were identified and 4,902 protein groups were quantified. The raw MS files and search/identification files obtained with MaxQuant have been deposited to the ProteomeXchange Consortium via the PRIDE partner repository with the dataset identifier PXD032156. (**F**) Summary of protein abundance of selected pan-endothelial and LEC-specific markers in HDLEC lysates. The sequence coverage (%), MS/MS counts per million spectra and relative abundance for each protein in HDLEC lysate are listed. Complete datasets are presented in the supplementary data (Supplemental file 4_Supplementary Table 1; Supplemental file 7_Supplementary Table Legend 1). (**G-H**) Pie charts reflecting the gene ontology (GO) analysis of protein class and subcellular localisation of members belong to quantified HDLEC proteome (Ashburner et al., 2000). GO terms were mapped using the Protein Annotation Through Evolutionary Relationship Evolutionary Relationships (PANTHER) classification system (Thomas et al., 2021; Mi et al., 2019). (**G**) Pie chart showing the relative distribution (percentage) of the different protein classes of quantified HDLEC proteome (mapping of 4890 out of 4902 HDLEC protein groups quantified in this study; 3628 classified gene IDs). (**H**) Pie chart showing the relative distribution (percentage) of the cellular compartments which quantified HDLEC proteome is associated with (mapping of 4890 out of 4902 HDLEC proteins quantified in this study; 7,608 hits of GO cellular compartment terms).

Immunoprecipitation (IP) of endogenous CLR from total primary HDLEC lysate with subsequent immunoblotting and quantification analysis demonstrated that both CLR forms (terminally and core glycosylated) were enriched with high efficiency (76.0±10%; Figures 2A, B). The fold-change difference of protein abundance between HDLEC total lysate and CLR IP eluate was also determined by the average LFQ intensity values of identified proteins (Supplemental file 2_Supplementary Figure 1; Supplemental file 3_Supplementary Figure Legend 1). The relative enrichment of CLR upon IP was also determined by MS-based protein abundance assigned by label-free quantitation (LFQ) intensity values (Figure 2C). Nine CLR peptides were identified by MS spectral counts using cultured *in vitro* HDLEC, seven more compared to a study used EC isolated from human skin tissue (ProteomeXchange Identifier PXD019909; Dyring-Andersen et al., 2020) (Figure 2D). The nine peptides detected in our study are located in the receptor’s N-terminus, third intracellular loop and the C-terminus, covering the 20% of the CLR sequence (Figure 2E).

**Figure 2.**
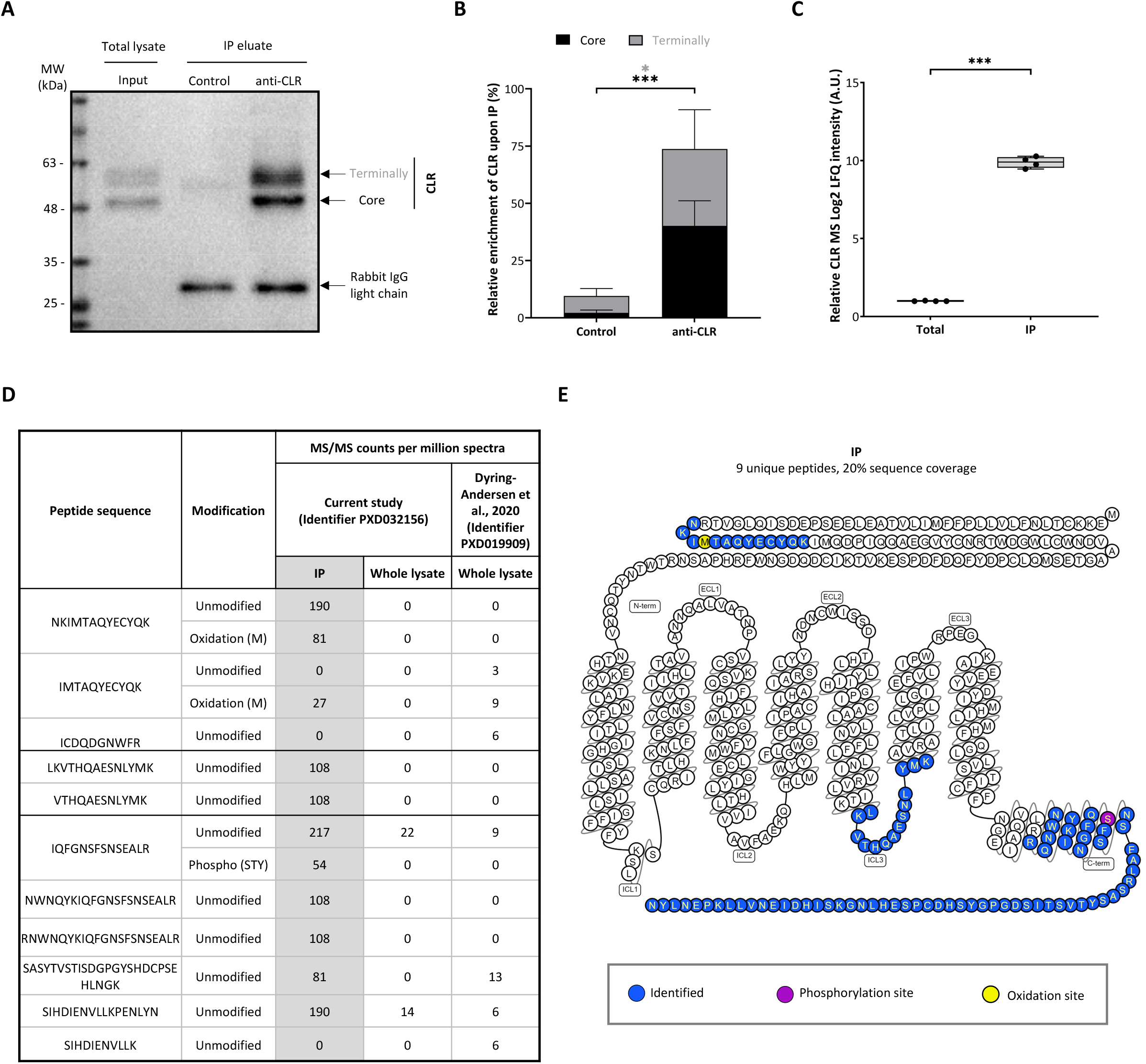
Detection and quantification of CLR in human dermal lymphatic endothelial cells upon immunoprecipitation and mass spectrometry analysis. (**A-B**) Calcitonin receptor-like receptor (CLR) immunoprecipitation (IP) from total human dermal lymphatic endothelial cell (HDLEC) lysates (input) was done using anti-CLR antibody/serum (Nikitenko et al., 2006). Non-immune serum was used as a negative control (n=4 independent experiments). (**A**) Detection and (**B**) relative quantification of CLR expression in eluate samples were done using immunoblotting (bars represent mean value ± SD; two-tailed unpaired t-test; *p<0.05, ***p<0.001). (**C**) Relative quantification of CLR expression in total lysate and IP samples, as based on the log_2_ label-free quantitation (LFQ) intensity acquired by nano-liquid chromatography-tandem mass spectrometry (nLC-MS/MS). (**D**) List of CLR peptides and their modifications identified by peptide to MS spectra counts in current study, using CLR IP eluate or whole cell lysates of *in vitro* cultured HDLEC, compared to endothelial cell isolated from skin tissue study (ProteomeXchange Identifier PXD019909; Dyring-Andersen et al., 2020). (**F**) Snake plot demonstrating CLR peptides identified in HDLEC in current study in HDLEC, was generated by using GPCRdb (Pándy-Szekeres et al., 2017).

Quantitative label-free nLC-MS/MS analysis of CLR co-IP proteome revealed enrichment in the expression of 46 protein identifiers in HDLEC (all novel), when a cut-off value of mean log_2_ LFQ intensity difference ≥2.25 (+1 SD) and FDR-adjusted *p*-value ≤0.01 compared to control (non-immune serum) was used (Figure 3A). Differential fold change enrichment of CLR IP proteome compared to total HDLEC lysate proteome was revealed by analysis of identified peptides count, LFQ intensity, sequence coverage and count of peptide spectra matches per million spectra (Figures 3B, C). From 46 novel CLR interactors, 11 were selected for antibody validation by immunofluorescence (IF) analysis using primary mouse monoclonal antibodies and paraformaldehyde (PFA) fixation which are essential for performing *in-situ* proximity ligation assay (PLA). IF analysis revealed detectable expression of six proteins, calcium/calmodulin dependent protein kinase II delta (*CAMK2D*), nucleoporin 93 (*NUP93*), serine/threonine-protein kinase MRCK beta (*CDC42BPB* or *MRCKB*), protein ERGIC-53 (*LMAN1*), valosin containing protein (*VCP*) and aconitase 1 (*ACO1*) in HDLEC (Supplemental file 2_Supplementary Figure 2; Supplemental file 3_Supplementary Figure Legend 2). These six antibodies and in-house rabbit anti-human CLR serum (Nikitenko et al., 2006) were used for PLA, which demonstrated that CLR directly interacts with CAMK2D, NUP93, MRCKB, ACO1 and VCP (Figures 3D, E). Quantified PLA signals were in accordance with the expression levels of these members of CLR in the co-IP proteome detected by MS (Figure 3C).

**Figure 3.**
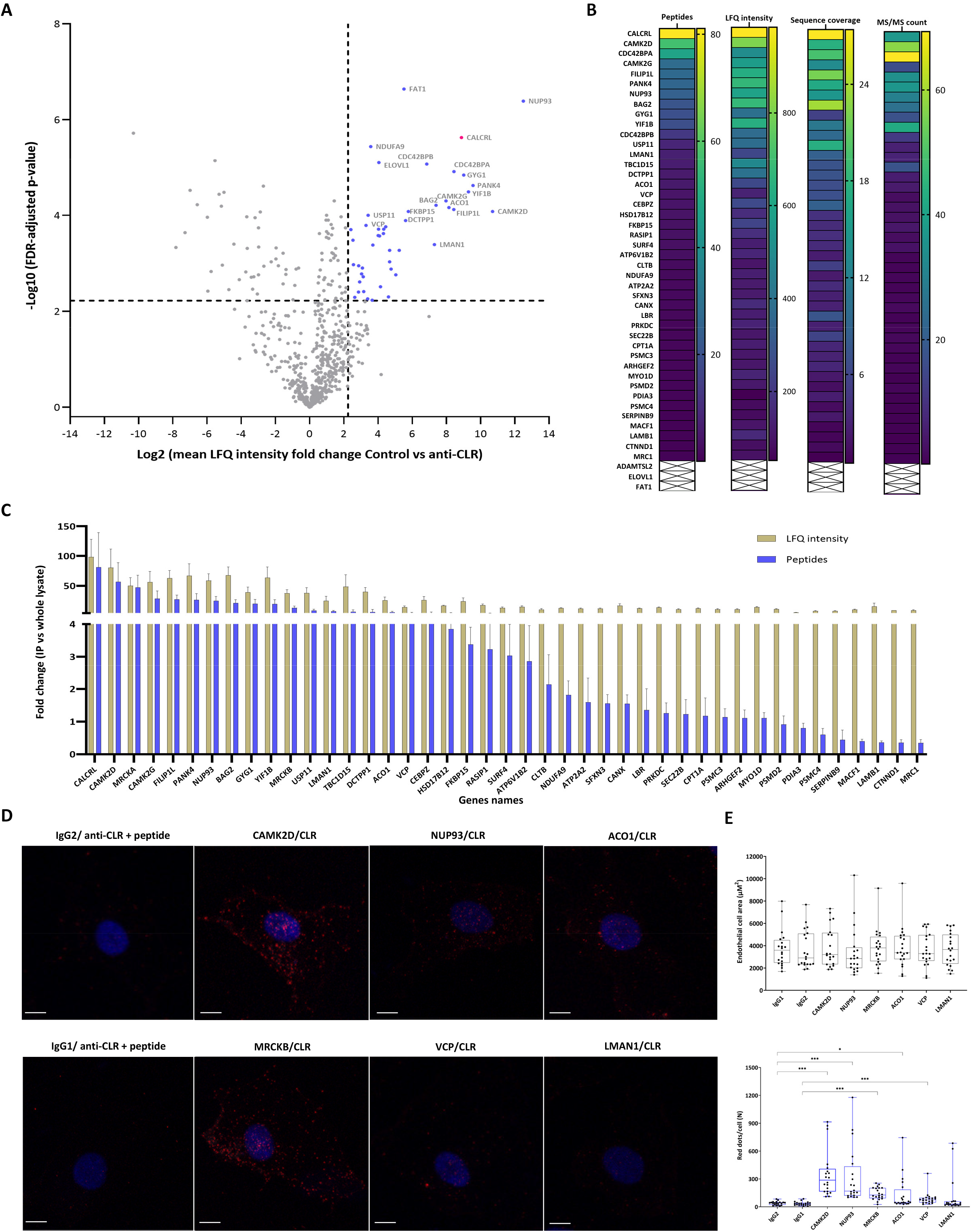
Identification and quantification of protein interaction network of endogenous CLR in human dermal lymphatic endothelial cells. (**A-C**) Proteins co-immunoprecipitated (co-IP) with calcitonin receptor-like receptor (CLR) from human dermal lymphatic endothelial cell (HDLEC) by using label-free nano-liquid chromatography-tandem mass spectrometry (nLC-MS/MS). (**A**) Volcano plot showing the log_2_ difference in protein enrichment of CLR co-IP proteome (anti-CLR serum LN-1436 against non-immune serum; please see Materials and Methods for full details) versus the minus log_10_ false discovery rate (FDR)-adjusted p-value (n=4; two-tailed student’s t-test). A permutation-based FDR of 0.01 was used for multiple hypothesis testing correction. Proteins were considered potential CLR binding partners if they showed a mean log_2_ label-free quantitation (LFQ) intensity difference ≥2.25 (+1SD of FDR-adjusted p-value population) and FDR-adjusted p-value≥2.23 in IP eluate samples using anti-CLR serum compared to non-immune serum as a control (46 protein identifiers). Complete datasets are presented in the supplementary data (Supplemental file 5_Supplementary Table 2; Supplemental file 7_Supplementary Table Legend 2). (**B**) Heatmaps of fold-enrichment of potential CLR interacting partners compared to whole HDLEC lysate was quantified based on identified peptides, LFQ intensity, sequence coverage and peptide to spectra match per million (MS/MS count) spectra. (**C**) Bar chart showing the fold-enrichment of identified members of CLR co-IP proteome compared to whole HDLEC lysate, based on identified peptides and LFQ intensity (10^-1^). (D-E) *In-situ* proximity ligation assay (PLA) was performed to detect and quantify CLR co-IP proteins in HDLECs. (**D**) Representative images of PLA signal (red dots) for 6 selected signature CLR co-IP proteins compared to their controls are shown. Nuclei were counterstained with DAPI (blue colour). Scale bars represent 10 μm. (**E**) Top- Box plot overlaid with dot plot represents the quantitation analysis of area of HDLEC used for PLA signal quantification (n=10 cells per group; mean value ± std; Two-way ANOVA). (**E**) Bottom- Box plot overlaid with dot plot represents the quantification analysis of PLA signal (dots/cell; n=10 cells per group; mean value ± SD; unpaired t-test or Mann-Whitney test; **P<0.01; ***p<0.001).

Gene Ontology cellular component (complete) and PANTHER protein class analyses demonstrated the relative distribution of the protein classes (Figure 4A) along with predicted subcellular localisation (Figure 4B) CLR binding partners in HDLEC. In this first study of its kind, we have identified a cohort of 46 novel protein binding partners of CLR which are endogenously expressed in HDLEC. Mapping of the identified CLR signature interactome revealed that immunoprecipitated core- and terminally- glycosylated CLR forms (Figure 1D) interact with a much larger network of proteins than previously thought (Figure 4C; representation of data from 4A and B). CLR interactome includes enzymes (such as kinases), transcriptional and signal transducers and modulators, while it is associated with cellular compartments where core- and terminally- glycosylated CLR is known to be localised, including Golgi apparatus, endoplasmic reticulum, intracellular vesicles and cell surface (Figure 4C).

**Figure 4.**
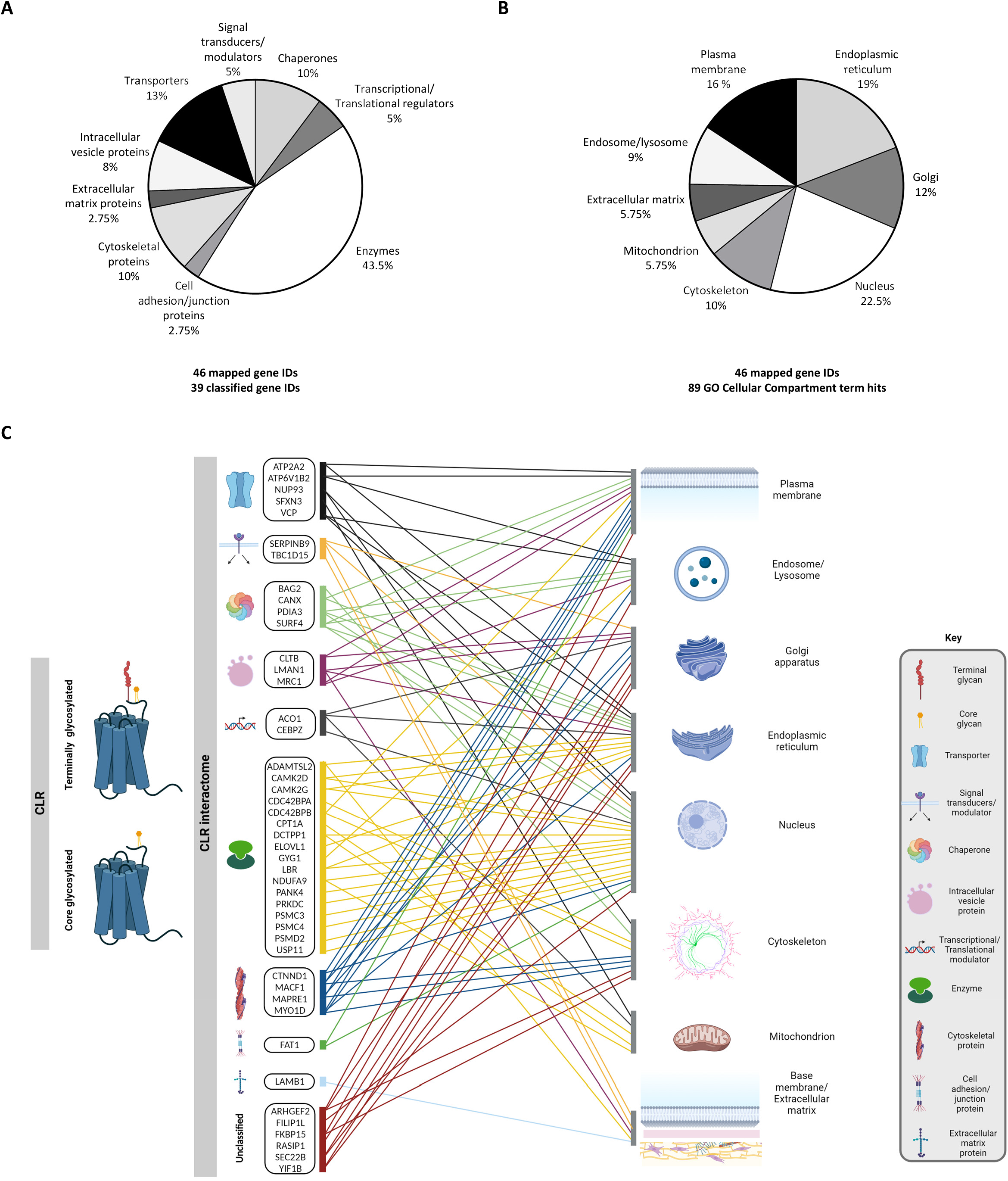
Classification and sub-cellular localisation of proteins belonging to CLR interactome in *in vitro* cultured human dermal lymphatic endothelial cells. (**A-B**) Pie charts reflecting the gene ontology (GO) analysis of protein class and subcellular localisation of members belong to calcitonin receptor-like receptor (CLR) interactome in human dermal lymphatic endothelial cells (HDLEC) (Ashburner et al., 2000). GO terms were mapped using the Protein Annotation Through Evolutionary Relationship Evolutionary Relationships (PANTHER) classification system (Thomas et al., 2021; Mi et al., 2019). Complete datasets are presented in the supplementary data (Supplemental file 6_Supplementary Table 3; Supplemental file 7_Supplementary Table Legend 3). (**A**) Pie chart showing the relative distribution (percentage) of the different protein classes of proteins belong to CLR interactome (39 classified gene IDs mapped for 46 gene identifiers). (**B**) Pie chart showing the relative distribution (percentage) of the cellular distribution of intracellular component the proteins interacting with CLR belong to (mapping of 89 hits GO cellular compartment terms for 46 gene identifiers). (**C**) Representation of data classification and predicted sub-cellular localisation of CLR interactome (from 4A and B). Core and terminally glycosylated forms of CLR, are shown on the left. The protein classes are represented by specific icons (Key). Each protein is connected (lines) with its relevant cellular component (defined by Gene Ontology classification; Ashburner et al., 2000; Mi et al., 2019), reflecting its predicted subcellular localisation. The scheme created with BioRender.

## Discussion

Proteins interacting with GPCRs ultimately define their functional properties and pharmacology in the cells and tissues expressing them. In this study, by combining co- immunoprecipitation, nLC-MS/MS and *in situ* PLA, we identified a novel group of proteins interacting with endogenous CLR in primary HDLEC cultured *in vitro*. From a total of 4,902 proteins detected in HDLEC lysates, we discovered 46 proteins which co- immunoprecipitated with core- and terminally-glycosylated CLR forms. These novel findings generate a platform for dissecting the effects of individual proteins from CLR interactome on the regulation and function of this GPCR in primary cells beyond HDLEC. Importantly, this knowledge will promote further fundamental and preclinical studies in human tissues and various pathologies ranging from cardiovascular disease and migraine to lymphoedema and cancer, where CLR roles have been implicated (Hoopes et al., 2012; Mackie et al., 2018; Nikitenko, Leek, et al., 2013; Nikitenko, Shimosawa, et al., 2013).

Our large-scale label-free quantitative MS-based proteomic study resulted in the acquisition of a more comprehensive (almost nine-fold increase) proteomic profile in primary HDLEC (∼5000 identified proteins) when compared to an analogous study, where Data Dependent Acquisition (DDA) mode was also used, and 561 proteins were detected (Roesli et al., 2008). In our study, two CLR peptides (6.7% amino acid sequence coverage; 47 MS/MS counts per million spectra) were identified in HDLEC lysates by nLC-MS/MS and CLR abundance was quantified amongst LEC markers, including LYVE1 (two peptides; 5.6% amino acid sequence coverage; 41 peptides to spectra matches per million spectra). This is less than in a study which identified six CLR peptides (17.1% amino acid sequence coverage) in the total cell lysate of primary EC isolated from human skin tissue (Dyring-Andersen et al., 2020), presumably due to abundant CLR expression in dermal blood vessel EC (Nikitenko, Shimosawa et al., 2013). In the current study, two forms of endogenous CLR (core- and terminally-glycosylated) were successfully immunoprecipitated for the first time and with high efficiency. The enrichment of endogenous CLR using IP enabled the detection of nine CLR peptides (20% amino acid sequence coverage), which is to our knowledge, the highest coverage for this GPCR identified in a single study to date. Our approach, therefore, led to the efficient isolation of CLR endogenously expressed in primary HDLEC, providing the capability to discover proteins which interact with it.

Intriguingly, CLR interactors previously identified in models of ectopic expression of tagged receptors (Hilairet, Bélanger, et al., 2001; McLatchie et al., 1998), such as RAMPs, clathrin and β-arrestins were not identified in our study. This is perhaps unsurprising, as for RAMPs, only one peptide (8% amino acid sequence coverage) of RAMP2 was previously identified in a total proteome from primary EC isolated from human skin tissue (Dyring-Andersen et al., 2020) and none from HDLEC in our study, highlighting the potential difficulty in detecting these accessory proteins by MS when they are endogenously expressed. Furthermore, clathrin heavy and light chains, along with β-arrestin 1, were only identified in the HDLEC total lysate, but not the CLR IP proteome. According to previous reports, ectopically expressed tagged CLR-RAMP1 complex internalises with β-arrestins upon 30 minutes stimulation of HEK293 cells with CGRP (Hilairet, Bélanger, et al., 2001). The absence of clathrin and β-arrestins in CLR interactome in HDLEC could be due to basal or steady-state (i.e. not stimulated with agonist) conditions used herein. These observations suggest that discovered by us proteins that associate with two CLR forms (core- and terminally- glycosylated) in HDLEC could represent a “steady-state interactome” of this GPCR.

In our study, 46 interactors of CLR were identified, and the direct interaction with the GPCR was confirmed for 5 of them using *in situ* PLA assay. Some proteins which co-immunoprecipitated with CLR have been previously shown to interact with and affect the trafficking and signalling of other GPCRs in different cell types (Roy et al., 2013; Semesta et al., 2020). More specifically, a cohort of CLR-interacting proteins (VCP, PSMD2, CANX and SEC61A1) has been shown to interact with the beta-2 adrenergic receptor (β2-AR) ectopically expressed in HEK293 cells, with VCP and NUP93 affecting the trafficking of this GPCR (Fan et al., 2005; Roy et al., 2013; Semesta et al., 2020). These studies also suggest that VCP is involved in the polyubiquitination of β2-AR in ER membranes and that NUP93 is required for both proper export to the plasma membrane and ligand-induced internalisation of this GPCR. NUP93 knockout has been associated with a significant reduction in total cAMP upon agonist-induced stimulation of β2-AR (Semesta et al., 2020). Furthermore, CAMK2G mediates phosphorylation of focal adhesion kinase upon agonist-activation of various GPCRs, such as bombesin, vasopressin, or bradykinin (Fan et al., 2005). Since members of the CLR interactome have the capacity to affect the function of other GPCRs in a pluripotent manner, it could not be excluded that CLR properties and signalling in HDLEC could also be regulated by them in a similar fashion.

To our knowledge, there are no other studies reporting the functional roles of the CLR-interacting proteins identified in our study in HDLEC or the lymphatic system. *In vivo* and *in vitro* studies have demonstrated that CLR and its agonists play important roles in dermal lymphatic system development and homeostasis (Chen et al., 2011; Davis et al., 2017; Hoopes et al., 2012; Jin et al., 2008; Nikitenko, Shimosawa et al., 2013; Tanaka et al., 2016; Tsuchiya et al., 2010). *CALCRL* mutation is lethal in human embryos (Mackie et al., 2018). However, to our knowledge, there are no known mutations in *CALCRL* gene in adults. We speculate/propose that inhibition of molecules that interact with CLR in HDLEC, as a key receptor in dermal lymphatic system function, would produce phenotypes resembling reduced CLR expression or affecting its function in these cells (Davis et al., 2017; Fritz-Six et al., 2008; Hoopes et al., 2012; Jin et al., 2008). To date, direct relevance of discovered in our study CLR binding partners to lymphatic biology and pathologies associated with lymphatic dysfunction, such as lymphoedema, has not been demonstrated. Therefore, further studies are needed to provide insights into how the CLR interactome could define its properties not only in lymphatic or other EC, but also in other cell types where this GPCR is expressed and involved in their biology.

The effects of several CLR-interacting proteins discovered in our study on other types of EC have been previously reported. In human umbilical vein EC (HUVEC), overexpression of FILIP1L inhibits cell proliferation and migration, while it is acting an apoptosis mediator (Kwon et al., 2008). In addition, RASIP1 required for the regulation of endothelial barriers function in these cells (Post et al., 2013). Furthermore, CAMK2D mediates enzyme- and growth factor-induced barrier and tube formation, while promotes cell migration and proliferation in HUVEC and human retinal microvascular endothelial cells (Wang et al., 2010; Ashraf et al., 2019). Altogether, the findings from studies using various EC types suggest that novel CLR interactors can possibly promote similar effects on HDLEC, facilitating the role of CLR and its agonists in proliferation, migration and monolayer permeability of these cells (Davis et al., 2017; Fritz-Six et al., 2008; Hoopes et al., 2012; Jin et al., 2008).

Apart from its abundant expression in HDLEC, CLR is also expressed in several other cell types, including human blood micro- and macro-vascular EC, vascular smooth muscle cells (VSMC), cardiomyocytes, neurons and cancer cells (e.g. in CCRCC and AML) in tissues and *in vitro* (Nikitenko et al., 2006; Nikitenko, Blucher et al., 2013; Hagner et al., 2002; Miller et al., 2016; Eftekhari et al., 2010; Angenendt et al., 2019; Gluexam et al., 2019). Importantly, this GPCR and its agonists are implicated in the pathogenesis and/or pathophysiology of several diseases, such as cardiovascular disease (dilated cardiomyopathy and heart failure), migraine and cancer (Kee et al., 2018; Nikitenko, Leek, et al., 2013; Scuteri et al., 2019; Voors et al., 2019; Zhang et al., 2018). The involvement of identified in our study CLR interactors in regulating function of various cell types, along with their role in pathology in many tissues has been also demonstrated. For example, CAMK2D plays a role in VSMC proliferation and migration, and is implicated in the pathogenesis of dilated cardiomyopathy and heart failure (Pfleiderer et al., 2004; Zhang et al., 2003; Cipolletta et al., 2010). In cardiomyocytes, CAMK2D is a key mediator of ER-stress responses and apoptosis, while ATP2A2 downregulation and GYG1 missense mutation have been associated with cardiomyopathy is adults (Zhu et al., 2007; Dally et al., 2010; Hedberg-Oldfors et al., 2017). LMAN1 regulates cell surface trafficking of neuroreceptors and CAMK2 is required for the induction of nerve injury-induced tactile allodynia in mouse hypothalamic and rat dorsal root ganglion neurons, respectively (Fu et al., 2019; Hasegawa et al., 2009). In CCRCC, upregulation of CLR expression in cancer cells is associated with survival outcome (Nikitenko, Leek et al., 2013), whilst PDIA3 expression increase cell proliferation and deubiquitinase USP11 interacts with tumour suppressor phosphatase and tensin homolog (PTEN) and diminishes its degradation (Liu, Wang et al., 2019; Zhang et al., 2019). In AML cells, CLR expression has been associated with more undifferentiated stage, which is linked to poor prognosis and resistance to therapy (Gluexam et al., 2019; Grandits & Wieser, 2021; Larrue et al., 2021; Nikitenko, Leek, et al., 2013; Angenendt et al., 2019), whilst CAMKG regulates cell proliferation, VCP promotes cell apoptosis via increased ubiquitination and high PANK4 expression has been associated with low survival outcome (Si et al., 2009; Szczęśniak et al., 2022; Liu, Cheng et al., 2019). Thus, our findings are likely to have a much wider, beyond lymphatic EC biology, impact on future studies on various cell and tissue types where the role of CLR and its agonists is implicated in normal and pathological conditions.

In summary, immunoprecipitation of endogenous CLR integrated with MS-based quantitative interactomics and *in situ* PLA led to the discovery of 46 novel binding partners for this GPCR in HDLECs. This integrative approach opens up the avenues for the discovery of CLR interactome and enhance understanding of the properties of this GPCR in a range of human cells and tissues, both in health and disease states, where it plays a significant role.

## Materials and methods

Full details of the methods are provided in the Supplemental file 1: Materials and methods.

## Declarations

### Ethics approval and consent to participate

Human dermal lymphatic endothelial cells (HDLEC) from a single female 29-years old donor were purchased commercially from PromoCell® (#C-12217; Lot number: 431Z012.3) and all ethical considerations (Human Tissue Act 2004) were met by the company, including donor’s consent.

### Consent for publication

Not applicable.

### Availability of data and materials’

The raw MS files and search/identification files obtained with MaxQuant have been deposited to the ProteomeXchange Consortium via the PRIDE partner repository with the dataset identifier PXD032156.

### Author contributions

DM, CE, HK and LLN contributed to study conception and design. DM and HK performed bioinformatic analysis. DM, SH and LLN performed immunofluorescence and image analysis. DM and LLN analysed and interpreted data. CE, AM, HK and LLN supervised study execution. DM and LLN drafted the manuscript. All authors reviewed the manuscript.

## Acknowledgements

Mr Chris Collins and Viper HPC Team, Professor Adrian Harris, Mr Eamon Faulkner and Mr Matthew Morfitt for their support.

