## Supplemental file 1 for "Novel protein interaction network of human calcitonin receptor-like receptor revealed by label-free quantitative proteomics"

**calcitonin receptor-like receptor**

**revealed by label-free quantitative proteomics**

Supplemental file 1_Materials and Methods

**Materials and Methods**

**Peptides and antibodies**

Primary and secondary antibodies were obtained from a range of manufacturers and used at dilutions and concentrations described below. Rabbit polyclonal anti-human CLR antibody in the form of serum (Nikitenko et al., 2006; LN-1436; 1:1000) was raised and characterised by us (Nikitenko et al., 2006). Primary mouse monoclonal CDC42BPA (#sc-374568); CDC42BPB (#sc-374597), ACO1 (#sc-166022), NUP93 (#sc-374400), LMAN1 (#sc-365158), CAMK2D (#sc-100362), GYG1 (#sc-271109), BAG2 (#sc-101216), DCTPP1 (#sc-398501) all from Santa Cruz Biotechnology and used at 3.0 μg/mL, TOP2B (#611492; 3.0 μg/mL); VCP (#612182; 3.0 μg/mL), CANX (#610523; 3.0 μg/mL), PECAM-1 (#555444; 1:100), CD144 (#555661; 1:150), CD107a (#555798; 1:200), EEA1 (#610456; 1:200), mouse IgG1 (#555746; 1:100) and mouse IgG2 (#555740; 1:100) all from BD Biosciences, PECAM-1 (#M0823; 1:50) and PDPN (#M3619; 1:50) were from Dako, mouse monoclonal beta-actin (#ab6276; 1:1000) was from Abcam, goat polyclonal PROX1 (#AF2727; 1:100), LYVE-1 (#AF2089; 1:100), rabbit IgG (#ab105C; 1:1000) and goat IgG (#ab108C; 1:200) were from R&D Systems, anti-rabbit IgG light chain (#NBP2-75935; 1:10000) was from Novus Biologicals. Secondary conjugated polyclonal donkey anti-rabbit IgG Alexa 594 (#A-21206; 1:600), anti-rabbit IgG Alexa 488 (#A-21207; 1:600), anti-mouse IgG Alexa 488 (#A-21202, 1:600), anti-goat IgG Alexa 488 (#A-11055, 1:600) and anti-goat IgG Alexa 594 (#A-11058, 1:600) all from Invitrogen, goat HRP anti-Mouse IgG (#P0447; 1:1000) and anti-rabbit IgG (#P0448; 1:1000) were from Dako.

**Cell culture, lysis and protein quantification**

*Ethical statement*

Human dermal lymphatic endothelial cells (HDLECs, passage 5) from a single 29 years old female donor were purchased commercially from PromoCell® were fully characterised and cultured as previously described (Nikitenko et al., 2006). Cells have been tested by the manufacturer for the absence of HIV-1, HIV-2, HBV, HCV, HTLV-1, HTLV-2 and microbial contaminants and by us for the lack of mycoplasma using EZ-PCR Mycoplasma Test kit (Biological Industries; #20-700-20) and the HyperLadder^TM^ 1kb (Bioline/Meridian Bioscience; # BIO-33026).

*Cell culture*

Cells were seeded onto a T-75 precoated flask and supplemented with PromoCell® EC growth medium MV2. Recombinant human VEGF-C (R&D Systems; #9199-VC; 7.5 ng/mL) was also added to the growth medium MV2. Cultures were incubated at 37°C in a 5% CO2 humidified atmosphere, and the medium was replaced every 24 hours. Cells were passaged 1:2 at confluence (~80%) by release with trypsin/EDTA (ethylenediaminetetraacetic acid) and cultured using the same method.

*Cell lysis*

Lysis was performed as previously described (Nikitenko et al., 2006). All steps were performed on ice. Cells were washed with ice-cold filtered PBS and homogenised using cell scrapers in radioimmunoprecipitation assay (RIPA) lysis buffer solution, in which protease and phosphatase inhibitor cocktails were included. Samples were processed in 1.5 ml tubes aspirating up and down and repeating three times with 10-minute intervals. Insoluble material was pelleted at 13, 000 g for 10 min at 4°C and supernatant was stored at -20°C prior to determination of total protein concentration.

*Quantification of total protein concentration*

Bicinchoninic acid (BCA) assay (Pierce™) was used for the determination of total protein concentration of cell lysates according to the manufacturer's instructions. The measurements were taken using a Tecan Infinite M200 Plate Reader and measuring absorbance at a wavelength of 562.0 nm.

**Immunoprecipitation (IP)**

A pre-cleaning step was performed prior immunoprecipitation, in which 2.0 μg of isotype control (rabbit non-immune serum) per 1,200 μg (2.0 μg/μl) of total protein were added to 4.0 mg of protein G magnetic beads (Invitrogen Dynabeads™ Protein G Immunoprecipitation Kit; # 10007D) and incubated by head-over-tail rotation for 30 minutes at 4 °C. For immunoprecipitation (Nikitenko et al., 2001), equal amounts of protein (600 μg; 2.0 μg/μl) were mixed with 2.7 μg of either anti-hCLR or isotype control serum and incubated by head-over-tail rotation for 90 minutes at 4 °C, in order to form the immune complex. Next, immune complexes were coupled to 5.0 mg of protein G magnetic beads and incubated by head-over-tail rotation for 60 minutes at 4 °C. The beads were washed three times using no detergent-based buffer (50 mM Tris, pH 7.4 and 150 mM NaCl, containing protease and phosphatase inhibitors) prior to subsequent elution steps. Immunoprecipitated samples were eluted by incubation at 55 °C for 25 minutes before SDS-PAGE and immunoblotting analysis. Alternatively, the washing buffer was removed and beads were stored at -80°C, before immune complexes were processed for on-bead tryptic digestion and mass-spectrometry analysis.

**SDS-PAGE and immunoblotting**

Protein lysates from cell lines and tissues were subjected to sodium dodecyl sulphate–polyacrylamide gel electrophoresis (SDS-PAGE) and immunoblotting as previously described (Nikitenko et al., 2006). Samples were electrophoretically separated on 10% polyacrylamide-based gel (acrylamide/methylene bisacrylamide solution at 37.5:1 ratio, 375 mM Tris pH 8.8, 0.1% SDS, 0.1% ammonium persulfate or APS and 0.04% tetramethylethylenediamine or TEMED) set with 5% stacking gel (acrylamide/methylene bisacrylamide solution at 37.5:1 ratio, 126 mM Tris pH 6.8, 0.1% SDS, 0.1% APS and 0.01% TEMED). Electrophoresis run using Tris running buffer at 100V for 120 minutes or until optimal resolution of proteins at 4 °C. Transfer to PVDF membrane was performed using Tris-based transfer buffer at 60V for three hours at 4 °C. The membranes were incubated in a blocking solution (5% non-fat milk in Tris-buffered saline containing 0.5% Tween-20 or TBS/T) for 60 minutes prior to primary antibody incubation. For primary antibody incubation, the membranes were incubated overnight in a blocking solution on a tube roller at 4°C. For secondary horseradish peroxidase (HRP)-conjugated antibody incubation, membranes were washed three times in 5-minute intervals using TBS/T, blocked as previously described and incubated at room temperature for 45 minutes when required. HRP activity was then detected using an enhanced chemiluminescence ECL kit. After detection, the membranes were stripped, reprobed, or stored at -20°C. Anti-human β-actin was used as control to monitor and confirm equal loading of total protein in samples.

*Protein detection and imaging*

Enhanced chemiluminescence (ECL) kit (BioRad) for the detection of HRP-conjugated antibodies was used. Imaging and densitometry were performed using Bio-Rad ChemiDoc XRS+ and Bio-Rad Image Lab 6.0 software respectively. Exposure times were varied and relied on the quality and intensity of the obtained signal.

**Immunofluorescence**

Immunofluorescence was used for HDLEC characterisation and antibody testing, as previously described (Nikitenko et al., 2006). HDLECs were sub-seeded in 8-well slide chambers (5000 cells per well) and upon reaching 80% confluency. For HDLEC characterisation and antibody testing, baseline conditions were used. Cells were washed and fixed using either 4% paraformaldehyde (PFA) or acetone/methanol (2:3 ratio) solutions. For 4% PFA fixation, the culture medium was removed from wells, which were washed with PBS prior to incubation with fixative for 7 minutes. Next, the PFA solution was removed and one wash with PBS was performed. PBS was added to fixed cells before their storage at 4°C. For fixation using acetone/methanol, the culture medium was removed, wells were washed once using PBS and incubated with fixative for 3 minutes. Next, the fixative was removed and the plates were left to air dry for 25 minutes, prior to their storage in cling film at -20°C. A pre-blocking step, using 10% donkey serum for 30 minutes at room temperature, was performed prior to primary antibody incubation. Primary antibodies were diluted at appropriate concentrations in 2% donkey serum and added to plates, wrapped in cling film and incubated overnight at 4°C. Incubation of human CLR serum with 10 μg/mL of the immunising peptide was used as CLR control staining (Nikitenko et al., 2006). Next, working on ice, wells were washed three times in 3-minute intervals with PBS before the incubation with secondary antibody in 2% donkey serum solutions. Incubation with secondary fluorophore-conjugated antibodies was performed under light protection at room temperature for 45 minutes. Next, the secondary antibody solution was removed and the wells were washed three times. When required, incubation with phalloidin (Invitrogen Alexa Fluor™ 635 Phalloidin; # A34054; 1:100) for 40 minutes at RT, was followed before mounting step with DAPI and imaging.

***In situ* Proximity Ligation Assay (PLA)**

For PLA with rabbit anti-human CLR, 11 mouse monoclonal antibodies, which have been routinely used in several studies were tested by IF at matching concentrations (3 μg/ml) and isotypes (IgG1 or IgG2). Antibody concentration was tested using SDS-PAGE prior to IF (data not shown). Cell cultured in 8-well chamber slides and washed once with PBS, fixed with 4% PFA for 7 min at 4 °C and briefly washed with PBS. Fixed cells were incubated with a blocking solution (10% donkey serum in 2% PBS-0.1% Triton X-100) for 30 minutes at RT. Next, cells were incubated with primary antibodies (in 2% donkey serum in PBS-0.1% Triton X-100) in a humidity chamber overnight at 4 °C. Incubation with 10 μg/mL of the immunising peptide used to raise anti-CLR serum was used as control of CLR staining. Next, fixed cells were washed in wash buffer (2x5 min in PBS-0.1%Triton X-100) at RT. Primary antibody solution was removed and fixed cells were washed 2x5 minutes in wash buffer at RT. PLUS and MINUS PLA probes were diluted 1:5 in wash buffer in a pre-heated humidity chamber at 37°C for 60 minutes. PLA probe solution was removed and slides were washed for 2x5 minutes in a wash buffer. Ligase was added at a 1:40 dilution in ligation buffer (1X) in high-purity water, and slides were incubated with the ligation solution in a pre-heated humidity chamber at 37°C for 30 minutes. Amplification reaction was conducted after 2x5 minutes washes with wash buffer, by incubating with amplification buffer (1X) and polymerase (1:80 dilution) solution in high purity water in a pre-heated humidity chamber at 37°C for 100 minutes. Incubation with phalloidin (Invitrogen Alexa Fluor™ 635 Phalloidin; # A34054; 1:100) for 40 minutes at RT, was followed after 2x10 washes in the washing buffer. Slides were dried in the dark at RT before mounting with DAPI for imaging.

**Microscopy and image analysis**

Confocal (ZEISS LSM 710) was used for HDLEC characterisation and antibody testing, and ImageJ/Fiji and Zen Blue 3.0 were used for image analysis. PLA fluorescence-stained cells were imaged using confocal microscopy (ZEISS LSM 710) with a 20x objective. All the PLA measurements were obtained using z-stacks of 5 images of 0.5 μm between each focal plane. Images were deconvolved and maximum projections were obtained in ZEN blue (Zeiss). Fiji/ImageJ was used for the semi-automatic quantitative assessment of PLA dots (Bertan et al., 2020). Images were taken with ROIs containing 5-8 cells. Based on the phalloidin staining, a F-actin mask (gaussian blur filter, subtract background, auto threshold “method=default”) was created to measure the cytoskeleton of each cell. The mask was used to count PLA signal (dots) using the Fiji/ImageJ option “find maxima” and possible off-target signals outside the cytoskeleton of each cell were excluded. PLA-dots were normalized based on cell area (μm^2^).

**Label-free mass spectrometry – sample preparation and data analysis**

For mass spectrometry analysis of immune complexes and whole cell lysates were subjected to proteolytic digestion, desalting and nano-liquid chromatography-tandem mass spectrometry (nLC-MS/MS). For on-bead-digestion of immunoprecipitated samples, beads were re-suspended using 4M urea in 20 mM HEPES (pH 8.0) solution. Immune complexes were incubated with 1.5 μg of LysC/Trypsin solution (Promega – PN: V5071, concentration 1.0 μg/μl) for 6 hours at 37°C. LysC is active at 4M urea. Next, bead slurry was diluted using HEPES and DTT (2.0 mM) solution in order to dilute urea to 1.0 M and DTT to 1.0 mM respectively. This dilution step will activate trypsin. Subsequently, samples were incubated overnight at 37°C. For alkylation and desalting*,* Iodoacetamide (5 mg/mL) was added to the samples and then incubated for 30 m in the dark. Samples were treated with 1 μl trifluoroacetic acid (TFA) to stop the digestion and desalted in C18 stagetips. For SP3 magnetic bead digestion of total cell lysates, a single‐pot solid‐phase‐enhanced sample preparation (SP3) method was used (Hughes et al., 2019). Protein samples were mixed with reconstitution buffer (50 mM HEPES, pH 8, 1% (wt/vol) SDS, 1% (vol/vol) Triton X-100, 1% (vol/vol) NP-40, 1% (vol/vol) Tween 20, 1% (wt/vol) deoxycholate, 5 mM EDTA, 50 mM NaCl, 1% (vol/vol) glycerol). Reducing agent stock (500 mM of DTT) was added to a final concentration of 5 mM DTT. Next, samples were heated using a Thermomixer at 60°C for 30 minutes, mixing at 1,000 rpm. Alkylating agent (chloroacetamide) was added to a final concentration of 20 mM and reaction was allowed to proceed for 30 minutes at room temperature. 100 μg of magnetic beads stock solution (50 μg/μl) were added to 10 μg of reduced and alkylated protein sample. 100% ethanol (to a final concentration of approx. 60%) was added and the solution was homogenised and the binding mixture was incubated in a Thermo-Mixer at 25 °C for 5 minutes at 1,000 rpm. The unbound supernatant was removed using a magnet and the beads were re-suspended using 80% ethanol solution. Next, the rinse was removed and 100 μl of digestion solution (100 mM ammonium bicarbonate pH 8.0 in water) containing 0.4 μg of trypsin per tube (for 1:50 trypsin to protein ratio) was added. Sonication for 1 minute on the ultrasonic water bath and incubation of fully reconstituted beads for 18 h at 37 °C in a Thermomixer at 1,000 rpm was followed. Digested samples were centrifuged at 20,000g for 1 minute and the supernatant was transferred into a fresh tube for each sample. Peptides were dried in a speed-vac and resuspended in 0.1% trifluoroacetic acid (TFA) solution (final concentration: 0.1% TFA).

For mass spectrometry data acquisition, peptides were analysed on a Q-Exactive mass spectrometer connected to an Ultimate Ultra3000 chromatography system incorporating an autosampler (both Thermo Scientific). 5 μl of the tryptic peptides, for each sample, was loaded on a homemade column (100 mm length, 75 μm inside diameter [i.d.]) packed with 1.9 μm ReprosilAQ C18 (Dr. Maisch, Germany) and separated by an increasing acetonitrile gradient, using a 40-min reverse-phase gradient (from 3%–32% Acetonitrile) at a flow rate of 250 nl/min. The mass spectrometer was operated in positive ion mode with a capillary temperature of 220 °C, with a potential of 2000 V applied to the column. Data were acquired with the mass spectrometer operating in automatic data-dependent switching mode, selecting the 12 most intense ions prior to tandem MS (MS/MS) analysis. Analysis of the raw data was performed using MaxQuant Linux version with the built-in Andromeda search engine (Cox & Mann, 2008; Tyanova, Temu, & Cox, 2016). The spectra were searched against the human UniProtKB/Swiss-Prot database version 06/2021 (canonical sequence). The MaxQuant default settings (including mass tolerance) were used. Specific settings: Trypsin as the protease (two missed cleavages); Carbamidomethylation (Cys) as the fixed modification; Oxidation (Met), Phosphorylation (Ser, Thr, Tyr) and N-terminal protein acetylation as variable modifications: The false discovery rate was 0.01 on both peptide and protein level and the minimum peptide length was set to seven amino acids. Quantification was done using the label free quantitation (LFQ) algorithm from MaxQuant (Cox et al., 2014). Mean LFQ intensities were calculated from technical-duplicate mass spectrometry runs for each of four biological replicates per experimental condition using MaxQuant and peptide and protein false discovery rates were set to 1%. LFQ intensity values were transformed to log2 values and sample and proteins quantified in fewer than 75% of all samples were excluded. Also, data was cleared of reversed hits, contaminants and “only identified by site. Missing values were imputed from a width-compressed, down-shifted normal distribution using Perseus version 1.6.7.0 (Tyanova, Temu, Sinitcyn, et al., 2016). The mean abundance of proteins was calculated using the ‘proteomic ruler’ package in Perseus (Wiśniewski et al., 2014).

**Gene Ontology (GO) enrichment analysis**

Functional profiling of the proteomic data was performed using the Gene Ontology resource from the GO Consortium server (Ashburner et al., 2000). Functional enrichment analysis of overrepresented ontology terms was performed with the GO Enrichment Analysis tool powered by PANTHER (Mi et al., 2019; Thomas et al., 2021). It allowed us to categorise the molecular function, biological process and cellular localisation of the unique proteins identified in this study.

**Statistical analysis**

IP-MS experiments were performed in quadruplicates. Results were deemed significant if P<0.05 and were denoted as: *P<0.05, **P<0.01 and ***P<0.001. For IP data, D'Agostino-Pearson test was used for normality. Two-tailed paired Student’s t-test was used for the quantification of immunoblotting data. For MS data, two-tailed paired Student’s t-tests with a permutation based on FDR-adjusted p-value=0.005 (FDR threshold of 5% applying 1000 randomizations) were conducted in Perseus (version 1.6.7.0). For analysis of PLA data n=10 cells per condition were used. D'Agostino-Pearson test was used for normality. Two-way ANOVA was used for the measurement of endothelial cell area (μm^2^) and the Mann-Whitney test for quantification of PLA signal (dots/cell). GraphPad Prism 8 software (San Diego, CA, USA) was used for statistical analysis unless stated otherwise. The different statistical tests used for individual experiments are described in figure legends.
