## Supplementary figures and images for "Novel protein interaction network of human calcitonin receptor-like receptor revealed by label-free quantitative proteomics"

### Supplemental file 2

##### Supplementary Figure 1

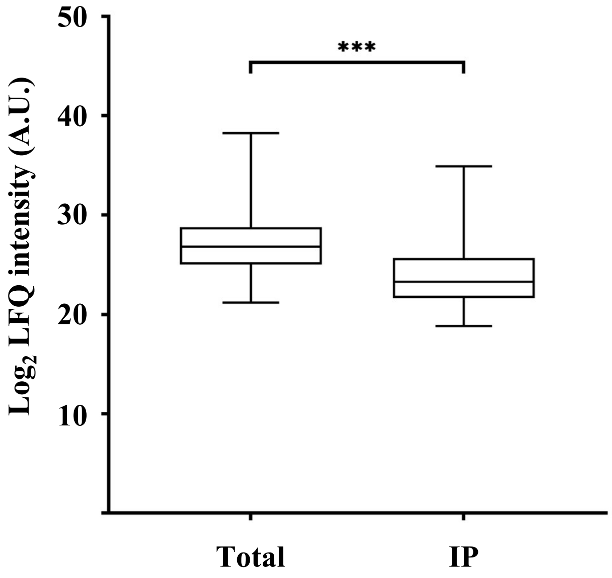

#### Supplementary Figure 2

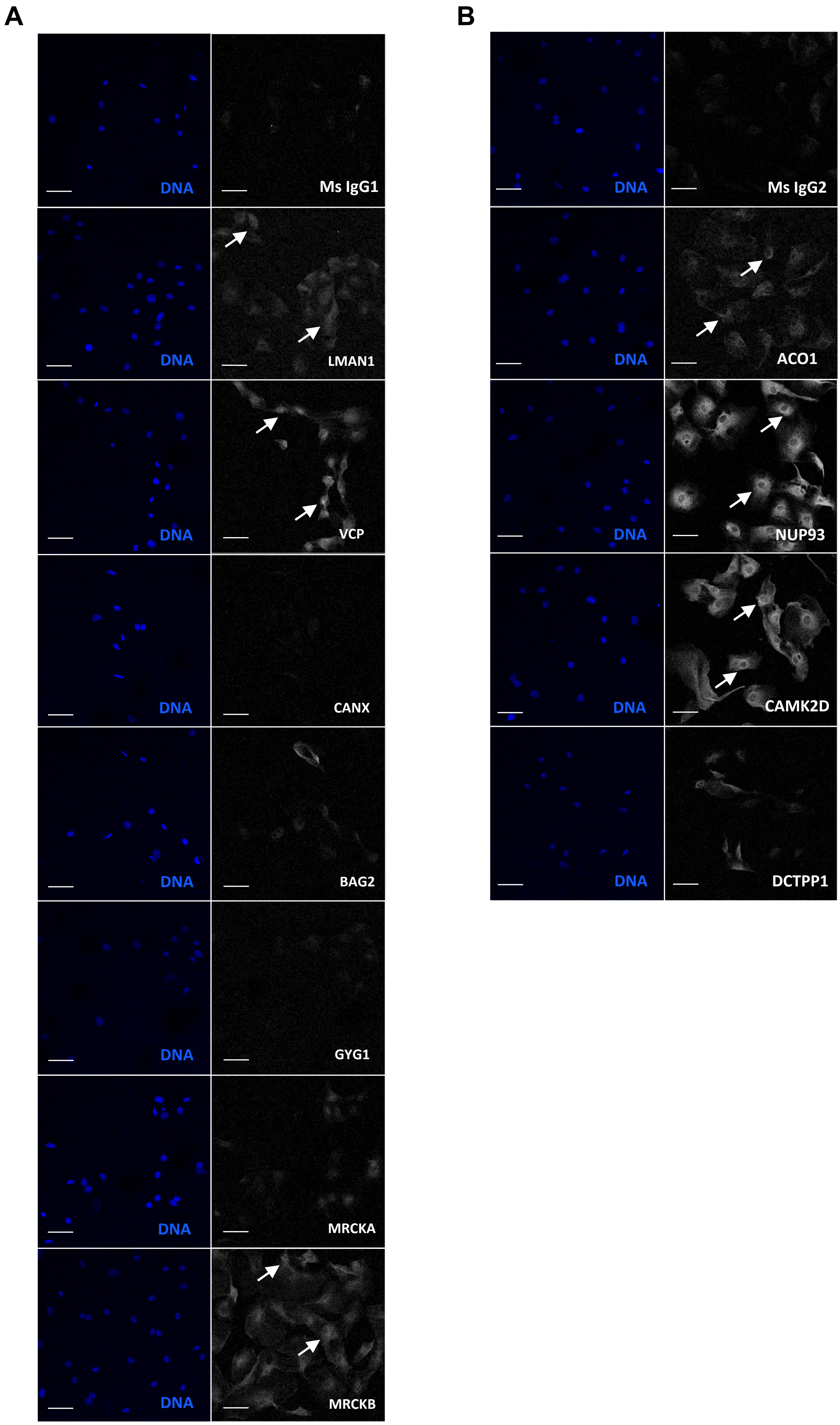
