## Supplemental file 3 for "Novel protein interaction network of human calcitonin receptor-like receptor revealed by label-free quantitative proteomics"

**calcitonin receptor-like receptor**

**revealed by label-free quantitative proteomics**

Supplemental file 3_Supplementary Figure Legends

**Supplementary Figure Legends**

**Supplementary Figure Legend 1**

**Fold-change difference of protein abundance between total lysate and CLR IP eluate in human dermal lymphatic endothelial cells.** Box plot is showing the protein abundance in total lysate (total) of human dermal lymphatic endothelial cells (HDLEC) and eluate samples upon calcitonin receptor-like receptor (CLR) immunoprecipitation (IP). Protein abundance was assigned by mean label-free quantitation (LFQ) intensity values acquired upon liquid chromatography tandem mass spectrometry. The LFQ values are plotted on a Log2(x) scale along the vertical axis. Statistical analysis was performed using D'Agostino-Pearson test (p<0.051; nonparametric distribution) and Mann-Whitney test (n=4 biological replicates; mean value ± SD; *** = p<0.001).

**Supplementary Figure Legend 2**

**Expression levels of 11 selected novel CLR interactors in human dermal lymphatic endothelial cells analysed by immunofluorescence.** Human dermal lymphatic endothelial cells (HDLEC) were cultured in vitro and fixed in 4% paraformaldehyde (PFA). See Materials and Methods for full experimental details. (A-B) Isotype controls (top images) IgG1 and IgG2 were used at matched concentration (3 μg/ml) to primary mouse monoclonal antibodies to analyse the expression levels of 11 selected proteins identified by label-free mass spectrometry as CLR interactors (genes LMAN1, VCP, CANX, BAG2, GYG1, MRCKA, MRCKB, ACO1, NUP93, CAMK2D and DCTPP1). Secondary Alexa Fluor® 488-conjugated antibody was used to reveal the signal (white pseudo colour; white arrows) for each protein. Nuclei were counterstained using DAPI (blue colour). Scale bars represent 20 μm.
