## Supplemental file 7 for "Novel protein interaction network of human calcitonin receptor-like receptor revealed by label-free quantitative proteomics"

**calcitonin receptor-like receptor**

**revealed by label-free quantitative proteomics**

Supplemental file 7_ Supplementary Table legends

**Supplementary Table legends**

**Supplementary Table Legend 1**

Identification and quantification of the proteome of primary *in-vitro* cultured human dermal lymphatic endothelial cells (HDLEC) by using label-free nano liquid chromatography-tandem mass spectrometry.

**Supplementary Table Legend 2**

Identification and quantification of proteins co-immunoprecipitated (co-IP) with calcitonin receptor-like receptor (CLR) from primary human dermal lymphatic endothelial cell (HDLEC) by using label-free nano-liquid chromatography-tandem mass spectrometry (nLC-MS/MS). The CLR interactome analysis was based on the mean log_2_ label-free quantitation (LFQ) intensity difference ≥2.25 (+1SD of FDR-adjusted p-value population) and FDR-adjusted p-value≥2.23 in immunoprecipitation (IP) eluate samples using anti-CLR serum compared to non-immune serum as a control.

**Supplementary Table Legend 3**

Classification and sub-cellular localisation of proteins co-immunoprecipitated (co-IP) with calcitonin receptor-like receptor (CLR) from primary human dermal lymphatic endothelial cell (HDLEC). Gene ontology (GO) terms of protein class and subcellular localisation were mapped using the Protein Annotation Through Evolutionary Relationship Evolutionary Relationships (PANTHER) classification system.
